# Photon-free (s)CMOS camera characterization for artifact reduction in high- and super-resolution microscopy

**DOI:** 10.1101/2021.04.16.440125

**Authors:** Robin Diekmann, Joran Deschamps, Yiming Li, Aline Tschanz, Maurice Kahnwald, Ulf Matti, Jonas Ries

**Affiliations:** Cell Biology and Biophysics Unit, European Molecular Biology Laboratory (EMBL), Heidelberg, Germany; LaVision Biotec GmbH, Bielefeld, Germany; Center for Systems Biology Dresden, Dresden, Germany; Department of Biomedical Engineering, Southern University of Science and Technology, Shenzhen, China; Collaboration for joint PhD degree between EMBL and Heidelberg University, Faculty of Biosciences, Heidelberg, Germany; Friedrich Miescher Institute for Biomedical Research, Basel, Switzerland

## Abstract

Modern implementations of widefield fluorescence microscopy often rely on sCMOS cameras, but this camera architecture inherently features pixel-to-pixel variations. Such variations lead to image artifacts and render quantitative image interpretation difficult. Although a variety of algorithmic corrections exists, they require a thorough characterization of the camera, which typically is not easy to access or perform. Here, we developed a fully automated pipeline for camera characterization based solely on thermally generated signal, and implemented it in the popular open-source software Micro-Manager and ImageJ/Fiji. Besides supplying the conventional camera maps of noise, offset and gain, our pipeline also gives access to dark current and thermal noise as functions of the exposure time. This allowed us to avoid structural bias in single-molecule localization microscopy (SMLM), which without correction is substantial even for scientific-grade, cooled cameras. In addition, our approach enables high-quality 3D super-resolution as well as live-cell time-lapse microscopy with cheap, industry-grade cameras. As our approach for camera characterization does not require any user interventions or additional hardware implementations, numerous correction algorithms demanding camera characterization become easily applicable.

---

(Scientific) complementary metal oxide semiconductor (sCMOS) cameras are increasingly popular for scientific imaging including fluorescence and super-resolution microscopy. For quantitative analysis of the images, pixelwise properties of the camera must be well characterized and accounted for in the analysis algorithm to avoid artifacts. This approach has been used to remove camera artifacts in both single molecule localization microscopy (SMLM)^1,2^ and diffraction-limited imaging^2–4^. Specific correction software is readily available^1,3–6^, but tools which can easily acquire the necessary data for pixel-dependent noise, offset, and photon response are still missing. Additionally, pixels feature individual dark current characteristics^5^, rendering both noise and offset functions of the camera exposure time, which is often neglected in characterization and correction algorithms. Consequently, a majority of (s)CMOS data is analyzed without explicit camera correction^7^. Industry-grade cameras approach the specifications of scientific-grade cameras and are increasingly used in the scientific community^8–15^. Especially for those cameras, a precise characterization and correction of the large pixelwise variability is indispensable.

Here, we developed a fully automated camera characterization pipeline, which determines pixel- and exposure time-dependent noise, offset and gain maps that are the basis for numerous camera correction algorithms. Our pipeline does not require any specific camera illumination, as it relies solely on dark current and associated thermal noise. In addition to gain, offset and noise maps, it also allows for the explicit consideration of dark current and thermal noise in the image reconstruction, which is of particular importance for long exposure times in SMLM or low light level live-cell imaging. We demonstrate that we can accurately characterize diverse (s)CMOS cameras and use the calibrations to avoid bias in 2D and 3D SMLM and in diffraction-limited imaging. Our camera characterization algorithm is implemented for the popular software packages Micro-Manager^16^ as well as ImageJ/Fiji^17^ and enables (s)CMOS specific corrections for the broad imaging community.

Camera characteristics are conventionally determined by evaluating mean and variance of the signal in each pixel over many images at several light levels^1^. The mean and variance of the signal with no light reaching the camera correspond to offset and read noise squared, respectively. Due to the stochastic and discrete nature of photon detection, the gain can be calculated as the ratio of the variance and mean signal at different light levels. Thermal excitation is an alternative source for excited electrons, resulting in dark current that increases both the offset and thermal noise and, with it, the overall noise. Accordingly, a calibration loses its validity when a different exposure time is used for imaging. This holds particularly true for long exposure times or uncooled cameras.

We turn this source of error to our advantage and use thermal excitations to fully characterize the camera without any light reaching the detector. This is possible as thermally excited electrons follow Poisson statistics just as photoelectrons. Photon-free camera characterization is based solely on dark current and thermal noise (Figure 1a), using the linear relation between exposure time and dark current to generate different signal levels (Figure 1b). Extrapolation to 0 ms exposure time gives the baseline (i.e. the offset free of thermal effects) as well as read noise (i.e. the noise free of thermal effects). Additionally, the explicit knowledge of the dark current and thermal noise as a function of exposure time now allows for computation of thermal effects at arbitrary exposure times (Figure 1c). For comparison, we used the traditional approach of varying light levels at a single frame exposure time of 10 ms (Figure 1d,e). Notably, the increased mean offset of 0.56 counts as compared to the photon free measurement equals the expected average dark current for 10 ms. For further verification, we calibrated an uncooled CMOS camera twice and found no considerable difference for all parameters (Supplementary Figure 1). We then compared the predictions of our approach to experimentally directly determined pixel-dependent offset and noise at different exposure times (Figure 1f). These comparisons showed high similarity, with average relative errors less than 0.4 % for the mean pixel values and 1.3 % for the noise. For the gain estimation (Figure 1e), we additionally compared our results with the single shot fluorescence method presented by Heintzmann et al.^18^ that is based on out-of-band information from diffraction limited fluorescence images. The relative deviation in the median gain by the different methods was below 3.4 %. To additionally consider variations in sensitivity (e.g. due to differences in quantum efficiency) between neighboring pixels, we optionally added the flat-fielding approach of Lin et al.^5^ and multiplied the flatfield map with the median gain to calculate the photon response map. However, for the cameras tested, the pixel-to-pixel variations in the flatfield map were very little (Supplementary Figure 2). Note that our approach does not correct for possible nonlinearities in the camera (Supplementary Figure 3), which would require more extensive characterization and correction routines^19^.

**Figure 1:**
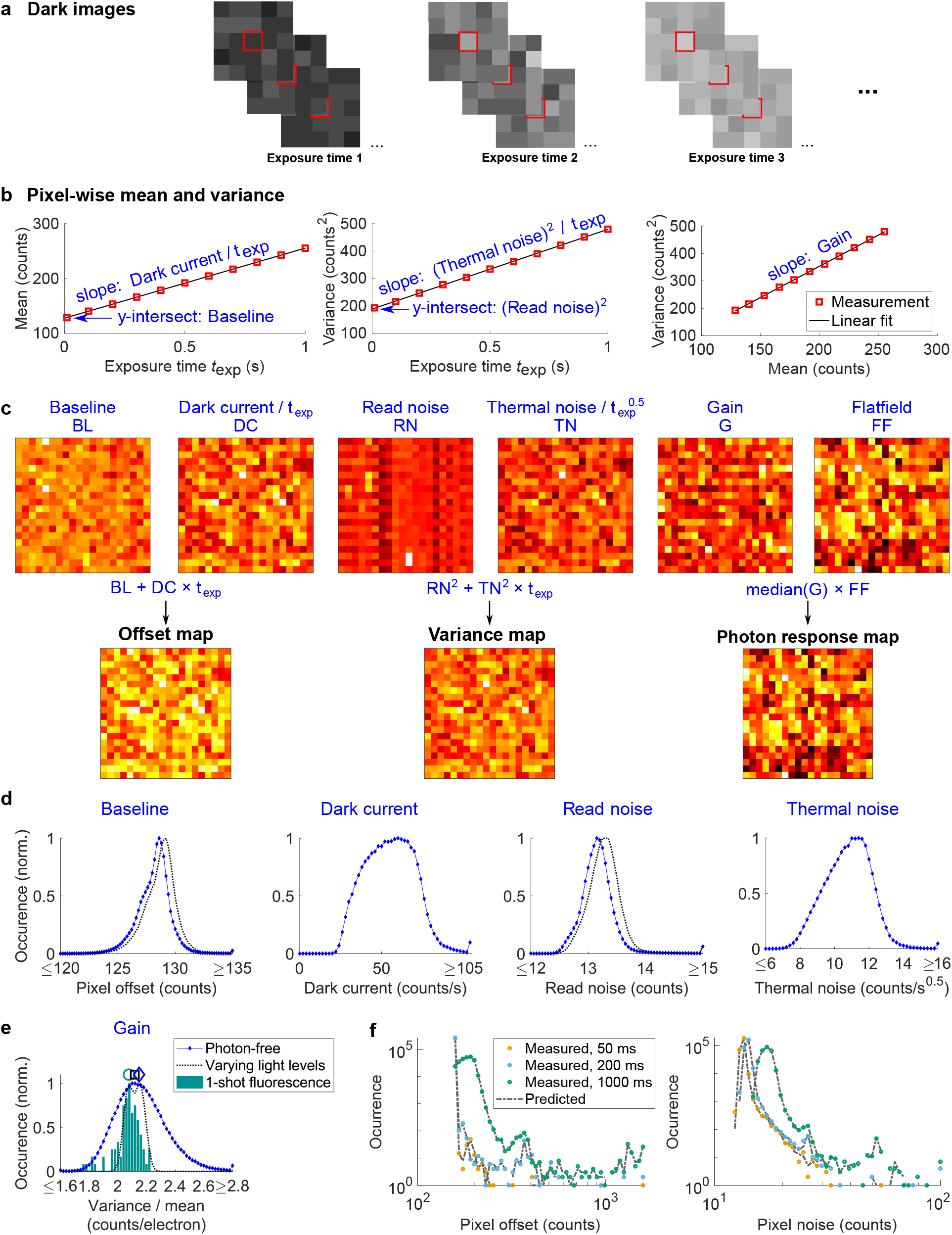
Automated camera characterization via thermally generated signal. **a**, A series of dark images is automatically recorded at different exposure times. For each pixel and exposure time the mean and variance of the signal is calculated. **b**, Result for one pixel of an uncooled CMOS camera. Dark current and thermal noise squared are proportional to the exposure time, so the temporal dependence can be determined from the slope of linear fits. The y-intersects of the fits reveal the baseline as well as the read noise squared, free from thermal effects. Since thermally generated signal follows Poisson statistics, the variance is proportional to the mean, with the proportionality factor corresponding to the pixel gain. **c**, Baseline, dark current, read noise, thermal noise and gain maps are calculated pixel-wise as in **b**. Optionally, we acquire a single bright image for flat-field correction. From these maps, we calculate the exposure-time dependent offset, variance and photon response maps. These maps can then be used as input for existing camera correction algorithms for images recorded at arbitrary exposure times. **d**, Histograms of pixel values obtained by photon-free characterization (blue curve) and traditional characterization of using varying light levels (black dashed curve). The traditional characterization overestimates baseline and read noise by the thermal effects for the corresponding exposure time. **e**, Distribution of the gain determined via different approaches (pixelwise histogram for the photon-free and varying light levels approaches, histogram of outcomes from multiple determinations of the mean gain from the 1-shot approach). Symbols above the curves indicate the medians (diamond for photon-free approach, square for vary light levels approach, circle for 1-shot fluorescence). **f**, Comparison of pixel offset and noise distributions at different single frame exposure times either predicted using the calculations shown in **c**, or directly determined from pixel-wise means and standard deviations.

**Figure 2:**
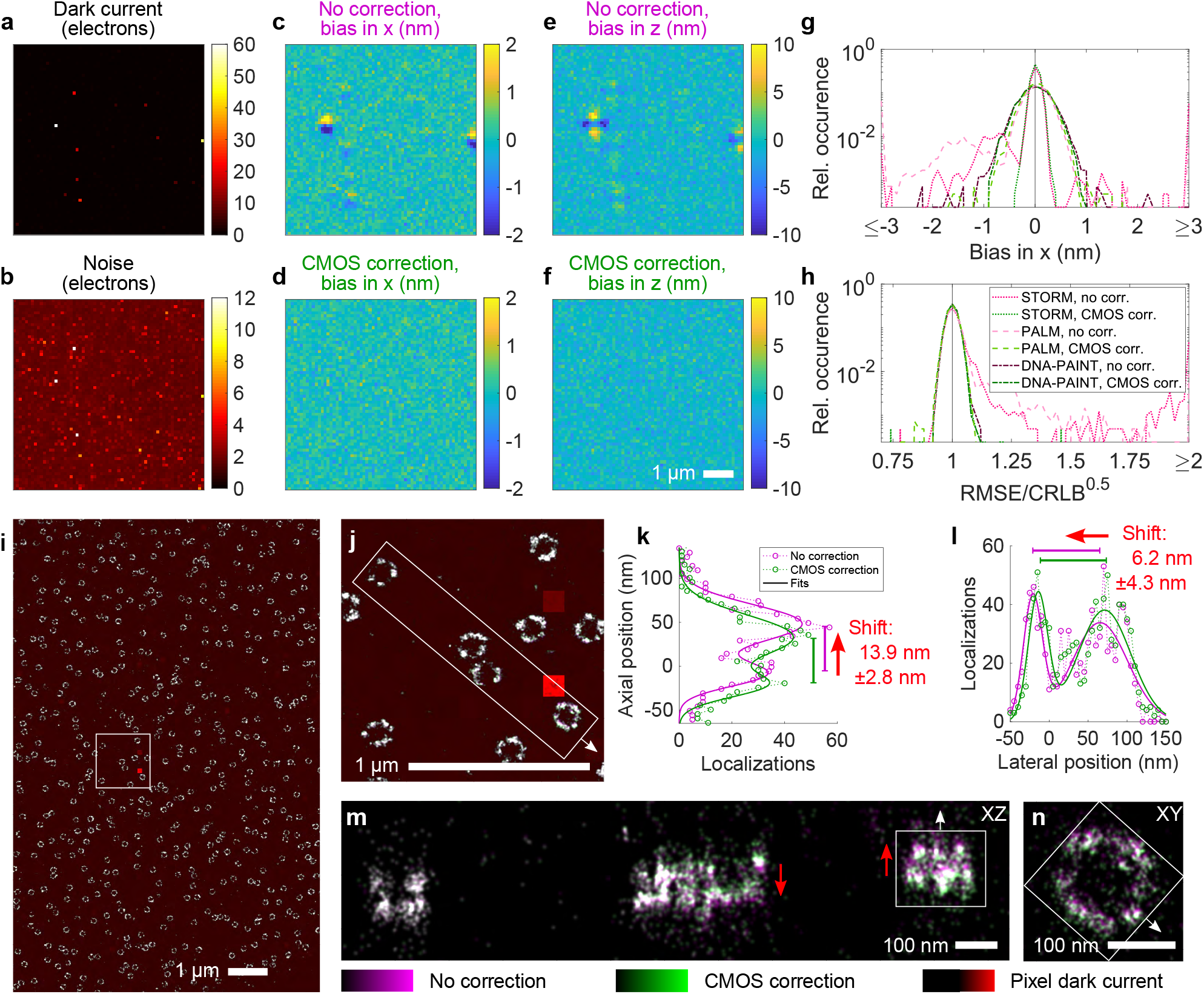
Camera calibration for thermal effects circumvents systematic fitting errors in SMLM. Maps of dark current (**a**) and noise (**b**) at 500 ms single frame exposure time for a scientific-grade CMOS (sCMOS) camera cooled to the manufacturer’s calibration setpoint of -10 °C. Simulations of 3D DNA-PAINT via astigmatism-based PSF shaping using this camera reveal a particular pattern in the localization bias close to pixels of high dark current, both laterally (**c**) and axially (**e**) when not applying (s)CMOS-specific fitting that corrects for pixel-wise effects including thermal effects. Explicit application of (s)CMOS specific fitting largely removes the bias for DNA-PAINT (**d**,**f**) as well as STORM and PALM (**g**) and restores the theoretically achievable root mean square error in the localizations (**h**). **i**, Experimental 3D DNA-PAINT data of the nucleoporin Nup96 in U2OS cells using the same cooled sCMOS camera. The image is rendered as an overlay of the pixel dark current map (red) and the SMLM reconstruction with no camera correction (magenta) and with CMOS correction (green). **j**, Zoom into boxed region in **i. k**,**l**, The structure of a nuclear pore complex (indicated by the boxes in **m**,**n**) becomes shifted in the vicinity of a pixel of high dark current, both in axial (**k**) and lateral (**l**) direction when neglecting individual pixel characteristics including thermal effects in the fitting pipeline. **m**, Axial view of the region indicated in **j**, also shown in Supplementary Video 1. **n**, Lateral close up of the nuclear pore complex indicated in **m**. As expected from the simulation, the shift features a high spatial dependence (**m**,**n**), which even changes its sign (indicated by the red arrows in **n**).

**Figure 3:**
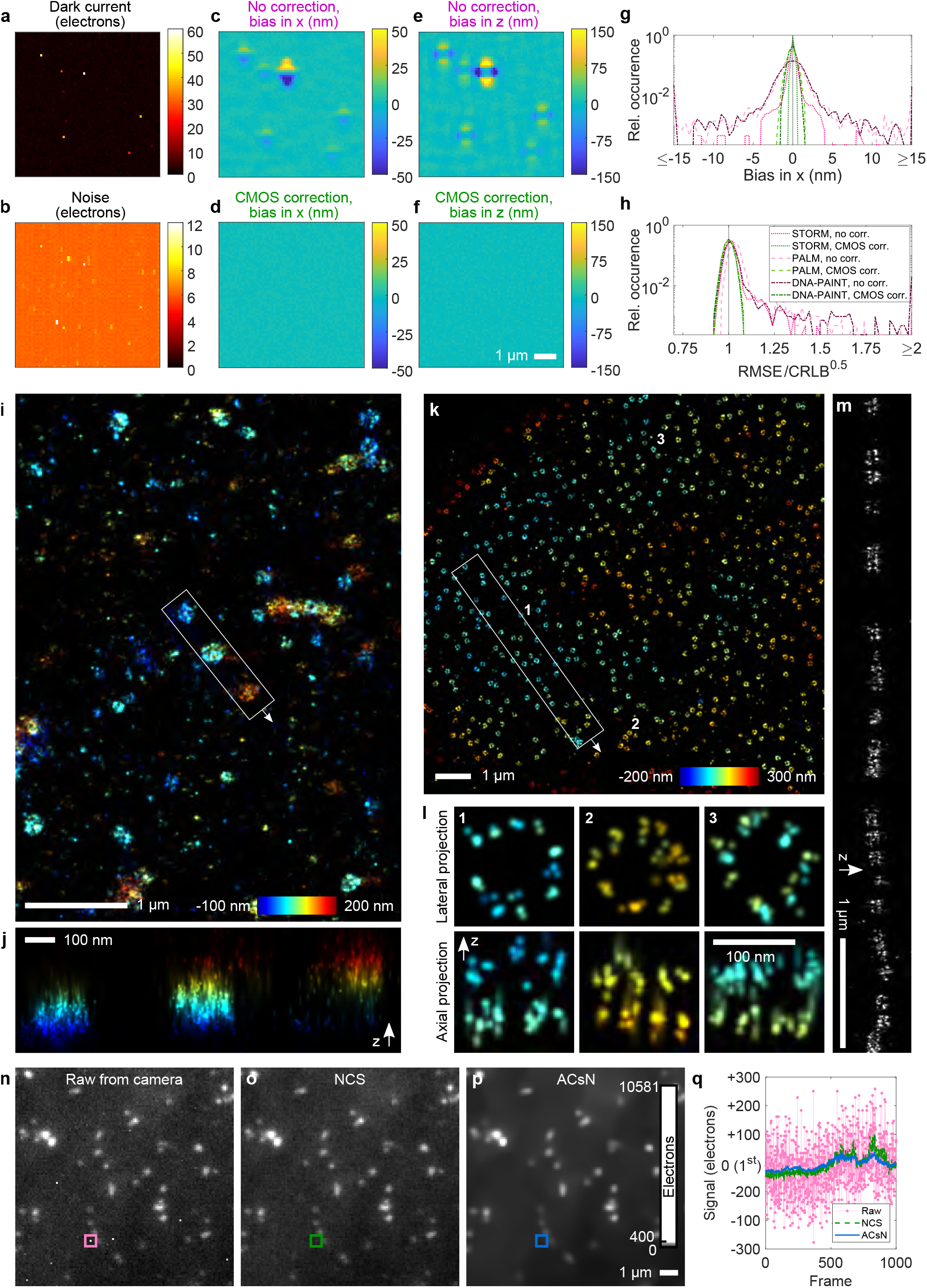
Camera calibration increases performance of an uncooled, industry-grade CMOS camera for SMLM and diffraction-limited fluorescence imaging. Maps of dark current (**a**) and noise (**b**) at 50 ms single frame exposure time for an uncooled, industry-grade CMOS camera (characteristics shown in Figure 1d,e). Simulations of astigmatism-based 3D PALM without explicit consideration of pixel-wise effects show a similar pattern in the localization bias, but of greater amplitude as compared to the cooled sCMOS camera (compare Figure 2), both laterally (**c**) and axially (**e**). Explicit application of CMOS-specific fitting largely removes the bias for PALM (**d**,**f**) as well as STORM and DNA-PAINT (**g**) and restores the theoretically achievable root mean square error in the localizations (**h**). **i**, Experimental 3D PALM data of clathrin tagged with mEOS3.2 in a U2OS cell using the same camera and applying CMOS-specific fitting including thermal effects. **j**, Axial view of region indicated in **i. k**, Experimental 3D STORM data of Nup107-SNAP, labeled with AF 647, in a U2OS cell using the same camera and applying CMOS-specific fitting including thermal effects. **l**, Gallery showing lateral and axial views on individual nuclear pore complexes indicated in **k. m**, The axial view on the region indicated in **k** shows two parallel lines from the nucleo- and cytoplasmic rings 57 nm apart. **n**, First frame of experimental raw data from time-lapse TIRF imaging of AP-2 tagged with eGFP in live U373 cells recorded at 1000 ms single frame exposure time. **o**, NCS corrected and, **p**, ACsN corrected frame. The entire time-lapse for **n-p** is shown in Supplementary Video 2. **q**, The noise of a pixel of high dark current is strongly reduced via both approaches after appropriate characterization of the camera including thermal effects. The signal has been offset-corrected by the pixel value of the first frame from the time-lapse.

Precise knowledge of the relevant (s)CMOS characteristics encoded in the offset, noise and photon response maps is crucial for accurate fitting in single molecule localization microscopy^1,5^ (SMLM). This holds especially true in the vicinity of hot pixels, which show increased offset and noise that strongly depend on the exposure time (Supplementary Figure 4). (s)CMOS-specific SMLM fitters^1,6^ which are based on maximum likelihood estimation (MLE)^20^ can achieve the theoretically achievable precision as given by the square root of the Cramer-Rao lower bound (CRLB). The camera maps generated by our approach (Figure 1) result in formally correct consideration of both dark current and thermal noise in (s)CMOS specific fitting (Supplementary Note 1) and we integrated the workflow into our SMLM software SMAP^21^.

To visualize the effect of (s)CMOS characteristics on SMLM, we simulated experiments of astigmatism-based^22^ 3D SMLM using measured maps of a latest generation, cooled scientific-grade CMOS camera (Fig 2a,b) and typical fluorophore parameters for traditional DNA point accumulation in nanoscale topography (DNA-PAINT)^23^ (i.e. long exposure time and high photon numbers), (direct) stochastic optical reconstruction microscopy (STORM)^24^ (i.e. short exposure time and medium photon numbers), and photoactivated localization microscopy (PALM)^25^ (i.e. short exposure time and low photon numbers) (Methods). When (s)CMOS specific fitting was not applied, regions close to pixels of high dark current show a high bias in the 3D localization coordinates (Figure 2c,e), even for DNA-PAINT for which sCMOS specific SMLM fitting is often neglected^26^. Application of sCMOS specific fitting largely removes the bias (Figure 2d,f,g), and restores the theoretically achievable root mean square error (Figure 2h) for all SMLM modalities.

To validate our simulation results with experimental data, we performed 3D DNA-PAINT^23^ using the same cooled scientific-grade CMOS camera. One might expect the bright fluorescence signal to be significantly higher than thermally generated signal. However, emitter dwell-times are in the 1 s regime, so dark current can play a pronounced role. Our experiments (Figure 2i,j) confirm the simulation results that the proximity of uncorrected high dark-current pixels leads to shifts in the localized coordinates. Although high dark-current pixels are relatively sparse on cooled, scientific-grade cameras, this bias locally misplaces structures in 3D (Figure 2k,l), easily exceeding nanometer localization precisions^26^. As expected from the simulations, the resulting distortion (Figure 2 k-n, Supplementary Video 1) is highly spatially dependent and changes its direction over only a few hundred nanometers. Consequently, even cooled sCMOS cameras should be characterized carefully and corrected for thermal effects for unbiased SMLM reconstructions. Besides DNA-PAINT, such characteristics (i.e. long exposure times and high photon numbers) are also relevant for STORM under resolution-optimized conditions^27^.

We next investigated if our approach can render uncooled, economic industry-grade CMOS cameras (Supplementary Figure 1) usable for high-quality 3D SMLM. Compared to sCMOS cameras, industry-grade CMOS cameras show higher dark current, higher noise and generally less uniform pixel properties^9^ (Figure 3a,b, all characteristics shown in Figure 1d-f). In simulations, these lead to an even larger bias in the localizations. Especially for PALM, local bias exceeded 50 nm laterally and 150 nm axially (Figure 3c,e). Again, consideration of pixel-dependent effects removes the bias (Figure 3d,f,g) and restores the theoretically achievable root mean square error (Figure 3h) for PALM, STORM and DNA-PAINT. Notably, applying general CMOS corrections but without explicit consideration of the exposure time retains bias (Supplementary Figure 5).

The potential of an uncooled, industry-grade CMOS camera becomes visible when we used it for 3D PALM^25^, i.e. SMLM with photoconvertible fluorescent proteins. PALM has lower signal levels as compared to STORM and DNA-PAINT, which is particularly challenging in presence of camera noise and for 3D SMLM. Following our photon-free camera characterization and applying the camera maps in CMOS-specific fitting, we could well resolve the 3D structure of clathrin coated pits by the same low-cost camera (Figure 3 i-j). 3D STORM^24^ (Figure 3k) on the nucleoporin Nup107 (ref. ^28^) clearly resolved individual corners of the nuclear pore complex (Figure 3l) in the lateral projection and parallel lines in the axial projection (Figure 3m) stemming from the nucleo- and cytoplasmic rings. This indicates a resolution better than 57 nm in the axial direction^29^, achieved with this uncooled, but properly characterized industry-grade camera. Thus, we could show that with our approach, low- cost cameras exhibit only slightly reduced performance compared to sCMOS cameras, an important development in light of their recent popularity for SMLM^8–10,12,14,15^.

Our CMOS characterization pipeline can also help in diffraction limited image restoration. Liu et al. have presented a noise correction algorithm (NCS)^3^ and Mandracchia et al. have presented an algorithm for automatic correction of sCMOS-related noise (ACsN)^4^. The aim of both approaches is to mitigate the effect of the (s)CMOS detector on wide-field images while preserving the characteristics of the fluorescence signal. Hence, both approaches rely on a proper camera characterization. NCS uses the three camera maps of offset, gain and noise. ACsN uses the two camera maps of offset and gain. We used an accordingly characterized, uncooled CMOS camera to image AP-2 in live U373 cells via total internal reflection fluorescence microscopy (TIRF)^30^ (Supplementary Videos 2). The raw data shows numerous pixels of strongly increased offset and noise (Figure 3n). Both NCS (Figure 3o) and ACsN (Figure 3p) removed noise and bias of such bad pixels (Figure 3q) and considerably increased the image quality.

Besides the conventional (s)CMOS corrections for pixel specific noise, gain and offset, our results indicate the benefit of characterizing and correcting for the effects of dark current and associated thermal noise in high- and super-resolution microscopy, both for uncooled and cooled cameras. To make our approach easily accessible for the imaging community, we implemented the *automated camera characterization via electron noise tool* (ACCeNT) in the popular open-source software Micro-Manager^16^ and ImageJ/Fiji^17^ (Supplementary Note 2). All relevant camera properties, including thermal effects, can be determined without user intervention and there is no need for additional hardware implementations. Thus, existing algorithms that demand proper camera characterizations, like the ones we used in this work, become applicable by a broad audience.

## Methods

### Camera calibration

For the photon-free calibration, all light to the camera chip was blocked by screwing a lid to the camera mount. Before starting the measurement, the camera was pre-run to give the detector time to thermally equilibrate either to the targeted cooling temperature (−10 °C as this was the manufacturer’s calibration setpoint) or warming up to the operating temperature in case of uncooled cameras. We acquired around 8,000 to 20,000 sets of typically 5 to 10 different exposure times. To maintain a constant average detector temperature, recording was performed in a nested manner, i.e. we changed the exposure time after each camera frame and then repeated acquisition of all exposure times.

Initially, we used custom-written scripts in Micro-Manager, Fiji and MATLAB for data acquisition and analysis, but later implemented the entire workflow into independent Micro-Manager and Fiji ACCeNT plugins (see next paragraph). After recording of raw data as described in the former paragraph, Fiji was used to process the TIFF stacks. For each exposure time and pixel, the mean value and standard deviation were calculated and saved as TIFF files: One after another, the stack corresponding to one exposure time was imported to Fiji and the “z-project” function was called with projection type “Average Intensity” for the mean value or projection type “Standard Deviation” for the standard deviation. The resulting TIFF files were imported into MATLAB for further processing. For each pixel linear functions were fitted using the polyfit function to (i) the mean value as a function of the exposure time to determine the baseline from the y-intersect and the dark current per time from the slope, (ii) the variance (i.e. the standard deviation squared) as a function of the exposure time to determine the read noise squared from the y-intersect and the thermal noise squared per time from the slope, and (iii) the variance as a function of the mean value to determine the gain (*i*.*e*. the conversion factor from electrons to ADU counts) from the slope. For each pixel, the exposure time dependent offset was calculated as the baseline plus the dark current per time multiplied by the single frame exposure time. For each pixel, the exposure time dependent noise squared was calculated as the read noise squared plus the thermal noise squared per time multiplied by the single frame exposure time. For each pixel, the photon response was calculated as the median of the gain map for all pixels multiplied by the pixel-wise value of the flatfield map. To find the flatfield map, we exposed the camera to a homogeneous illumination via ambient light and applied the algorithm presented by Lin et al.^5^.

We implemented the photon-free calibration workflow including the automated nested data acquisition, fitting of individual pixel properties and calculation of exposure time dependent camera maps as the ACCeNT plugin for Micro-Manager 2. Additionally, we implemented the fitting of individual pixel properties and calculation of exposure time dependent camera maps as an ACCeNT plugin for Fiji. The latter is intended to be used for processing of data acquired with software different from Micro-Manager 2, e.g. if the microscope is run using Micro-Manager 1.4 or the manufacturer’s software. For Micro-Manager 1.4 users, we provide a script for the automated nested data acquisition. We checked the consistency of all implementations against each other.

### Microscope

All data was acquired on a custom-built microscope as described in the following. Laser light was emitted from the single mode fiber output of a laser box (iChrome MLE, Toptica) (640 nm for excitation of Alexa Fluor 647 and Atto 655 and TetraSpeck beads, 561 nm for excitation of mEos and TetraSpeck beads, 488 nm for excitation of GFP and 405 nm for active photoswitching in STORM and PALM experiments) and collimated using an achromatic lens (either f = 50 mm for DNA-PAINT imaging using the industry-grade CMOS camera or f = 30 mm for live-cell experiments and all STORM and PALM experiments, all lenses from Thorlabs) or emitted from a fiber laser (F-04306-107, MPB Communications) (642 nm for excitation of Atto 655), filtered through an AOTF (AA Opto Electronic) and expanded using telescope of achromatic lenses (f = 50 mm and f = 100 mm, both Thorlabs). The collimated laser light was focused (f = 150 nm, Thorlabs) to the back focal plane of a TIRF objective lens (either 100x, NA 1.35 silicone oil, Olympus for DNA-PAINT using a cooled sCMOS camera, or 60x, NA 1.49 for all other experiments). Imaging the fiber output in 4f-configuration and mounting it to a linear stage (SLC2445me-4, Smaract) enabled image acquisitions in epi, HILO and TIRF illumination. The resulting projected pixel widths were 98 nm for an uncooled, industry-grade CMOS camera (µeye UI-3060CP-M-GL R, IDS), 58 nm for a different uncooled, industry-grade CMOS camera (Chameleon3 CM3-U3-50S5, FLIR), and 117 nm for a cooled, scientific-grade CMOS (sCMOS) camera (Edge 4.2 bi, PCO).

Fluorescence emission was separated from the laser excitation via a dichroic beamsplitter (zt405/488/561/640rpc, Chroma), further filtered (either bandpass 697/58, Semrock for DNA-PAINT using a cooled sCMOS camera; bandpass 700/100, Chroma plus notch filter 400-410/488/561/631-640, Semrock, for DNA-PANT using an uncooled, industry-grade CMOS camera; bandpass 676/37, Semrock for STORM; longpass 568, Semrock, plus bandpass 600/60, Chroma for PALM imaging; or 525/50, Semrock for live-cell GFP imaging) and focused onto the camera by a tube lens (either f = 100 mm, Thorlabs for DNA-PAINT using the cooled sCMOS camera; or f = 180 mm, Olympus for all other experiments). For 3D SMLM experiments via astigmatism-based PSF shaping, a cylindrical lens was placed before the camera (f = 2000 mm, CVI Laser Optics). An additional short pass filter (FESH750, Thorlabs) was used before the camera to block light from a focus lock laser. The focus lock laser (785 nm, Toptica) was coupled into the excitation beam path using an additional dichroic mirror, reflected off the coverslip-buffer interface of the sample, and its position was detected using a four-quadrant photodiode. The photodiode output was used to maintain the z-position of the objective lens constant with respect to the sample for active z-drift compensation.

The microscope hardware and data acquisition was handled via Micro-Manager 1.4.22 using custom-written software^31^. When imaging using an industry-grade CMOS camera, the excitation laser was run constantly. When imaging using the cooled, scientific-grade sCMOS camera, the excitation laser was triggered on during the common exposure of all lines of the camera. In all cases, the UV laser for active photoswitching in PALM and STORM experiments was triggered at the camera frame rate, but the pulse length was dynamically adjusted to aim for a constant number of active emitters per frame.

### Sample preparation

#### U2OS cells NUP107-SNAP for STORM

U2OS Nup107-SNAP samples stained with Alexa Fluor 647 for STORM imaging were prepared as previously described^29^.

#### U2OS cells clathrin mEOS3.2 for PALM

U2OS cells were seeded onto clean 24 mm round glass coverslips and grown in phenol-red free Dulbecco’s Modified Eagle Medium growth medium (DMEM, Gibco no. 11880-02; 1x MEM NEAA, Gibco no. 11140-035; 1x GlutaMAX, Gibco no. 35050-038; 10% [v/v] fetal bovine serum, Gibco no. 10270-106) (cell culture and seeding conditions described in ref. ^28^). Transient transfection with a clathrin-mEOS3.2 construct (Addgene 57452) was achieved using Lipofectamine^™^ 2000 reagent (Life Technologies) according to the manufacturer’s recommendations: DNA (1 ug) was mixed with OptiMEM I (50 uL), and Lipofectamin (3 uL) was mixed with OptiMEM I (50 uL). Both solutions were incubated for 3 min at room temperature, mixed together and incubated for additional 10 min at room temperature. After exchanging the culture medium with prewarmed OptiMEM I, the DNA-Lipofectamin solution (100 uL) was added dropwise to the seeded cells. After approximately 24 hours incubation (at 5% CO_2_, 37 °C), the medium was exchanged with fresh growth medium. After additional incubation for approximately 24 hours, cells were fixed for 20 min in 3% [w/v] paraformaldehyde in cytoskeleton buffer (CB; 10 mM MES pH 6.1, 150 mM NaCL, 5 mM EGTA, 5 mM D-glucose, 5 mM MgCL_2_, as described in ref. ^32^) at room temperature. The fixation process was stopped by incubation for 7 min in 0.1% [w/v] NaBH_4_ at room temperature. The sample was washed 3 times for 5 min in PBS.

#### U373 cells AP2-eGFP for live-cell TIRF

U373 cells stably expressing AP2-eGFP (generously provided by the Boulant lab, German Cancer Research Center (DKFZ), Heidelberg) were cultured in high glucose growth medium (DMEM, Gibco no. 11880-02; 1x MEM NEAA, Gibco no. 11140-035; 1x GlutaMAX, Gibco no. 35050-038; 10% [v/v] fetal bovine serum, Gibco no. 10270-106; 20% [w/v] glucose; 1x ZellShield, Minerva Labs) at 37°C and 5% CO_2_. Passaging was done every 2-3 days to maintain the cells at approximately 50% confluency.

#### U2OS cells immunostained for DNA-PAINT

U2OS cell immunostained for microtubules for DNA-PAINT imaging were prepared as previously described^6^ using anti beta-tubulin antibody (T5293; Sigma-Aldrich).

#### U2OS cells Nup96-eGFP for DNA-PAINT

Cells were seeded as previously described on high-precision 24 mm round glass coverslips^33^. In short, coverslips (No. 1.5H, catalog no. 117640, Marienfeld) were cleaned in a methanol:hydrochloric acid (50:50) mixture overnight before washing them repeatedly with ddH_2_O and drying them in a laminar flow hood. Before usage, clean coverslips were additionally irradiated with UV for 30 min.

U2OS Nup96-mEGFP cells were seeded onto the coverslips in such a density, that they reach a confluency of 50 to 70% on the day of fixation (typically 2 days after seeding). During this time, cells were grown in an incubation chamber providing 37 °C and 5% CO2 in growth medium (DMEM (catalog no. 11880-02, Gibco)) containing 1× MEM NEAA (catalog no. 11140-035, Gibco), 1× GlutaMAX (catalog no. 35050-038, Gibco) and 10% (v/v) fetal bovine serum (catalog no. 10270-106, Gibco). Finally, shortly before fixation, coverslips were rinsed twice with warm PBS.

Coverslips containing U2OS Nup96-mEGFP cells (catalog no. 300174, CLS Cell Line Service, Eppelheim, Germany) were first prefixed in 2.4% w/v formaldehyde (FA) in PBS for 40s before samples were incubated in 0.1% v/v Triton X-100 in PBS for 3 min. After washing samples twice for 5 min in PBS, fixation was completed in 2.4% w/v FA in PBS for 20 min. The sample was subsequently washed twice in PBS for 5 min each before remaining FA was quenched in 100 mM NH_4_Cl in PBS for 5 min and then washed twice in PBS for 5 min. Permeabilization was carried out in 0.2% v/v Triton X-100 in PBS and remaining permeabilization solution was washed away twice in PBS for 5 min each. Samples were blocked in 2% w/v BSA in PBS for 1 h, before coverslips were placed upside down onto a drop of primary antibody staining mix (rabbit anti-GFP, catalog no. 598, MBL International, diluted 1:250 in PBS containing 2% w/v BSA) overnight at 4 °C. Weakly and unbound primary antibodies were washed off thrice in PBS for 5 min each. Similarly, binding of anti-rabbit secondary i1 DNA-PAINT antibodies (homemade, kind gift of Ingmar Schoen, Royal College of Surgeons in Ireland) was achieved by placing the samples upside down onto a 1:100 dilution of the antibodies in PBS containing 2% w/v BSA for 1 h at RT. After washing thrice in PBS for 5 min each, a post-fixation was carried out in 2.4% w/v FA for 30 min. Samples were washed twice for 5 min in PBS and finally placed into a custom-made sample holder.

#### Fluorescent bead samples

100 nm sized TetrackSpeck beads (Thermo Fisher) were diluted 1:40 in 100 mM MgCl2 in H2O and incubated for 3 min on coverslips. Before imaging and PSF calibration via z-stacks, the bead solution was replaced by H2O.

### Data acquisition

PALM imaging was performed using an uncooled, industry-grade CMOS camera (µeye UI-3060CP-M-GL R, IDS). Fixed U2OS cells were imaged in buffer containing 95 % D2O and 50 mM Tris/HCl pH9. Raw data was acquired in HILO illumination at 561 nm and laser output powers of 20 mW to 50 mW. The single frame exposure time was set to 50 ms.

STORM imaging was performed using either an uncooled, industry-grade CMOS camera (µeye UI-3060CP-M-GL R, IDS) or a different uncooled, industry-grade CMOS camera (Chameleon3 CM3-U3-50S5, FLIR). Fixed U2OS cells were imaged in blinking buffer containing 50 mM Tris/HCl pH8, 10 mM NaCl, 10% (w/v) D-glucose, 500 µg/ml glucose oxidase, 40 µg/ml catalase, 143 mM BME and 2 mM COT. Raw data was acquired in HILO illumination at 640 nm and at a laser output power 70 mW. The single frame exposure time was set to 50 ms.

DNA-PAINT imaging of tubulin in U2OS cells was performed using an uncooled, industry-grade CMOS camera (µeye UI-3060CP-M-GL R, IDS). Fixed U2OS cells were imaged in buffer containing 500 mM NaCl, 1x PBS, 40 mM Tris/HCl pH8 and imager strands (I1 650, i.e. Atto655, Ultivue) at a concentration of about 500 pM. Raw data was aquired in HILO illumination at 640 nm and a laser output power of 70 mW. The single frame exposure time was set to 500 ms.

DNA-PAINT imaging of Nup96 in U2OS cells was performed using a cooled, scientific-grade CMOS (sCMOS) camera (Edge 4.2bi, PCO). Fixed U2OS cells were imaged in buffer containing 500 mM NaCl, 40 mM Tris/HCl pH8 and imager strands (I1 650, i.e. Atto 655, Ultivue) at a concentration of about 500 pM. Raw data was acquired in HILO illumination at 642 nm and a laser output power of 4.5 mW. The single frame exposure time was set to 500 ms.

Diffraction limited TIRF imaging of AP-2 in U373 cells was performed using an uncooled, industry-grade CMOS camera (µeye UI-3060CP-M-GL R, IDS). Live U373 cells were imaged at room temperature in growth medium. Raw data was acquired in shallow TIRF illumination at 488 nm and a laser output power of 0.1 mW. The single frame exposure time was set to 1000 ms.

### Image data analysis

SMLM data was fitted and analyzed as previously described^28^ using our custom-written, open-source superresolution microscopy analysis platform SMAP^21^ in MATLAB (Mathworks). The software is available at github.com/jries/SMAP. In case of (s)CMOS specific fitting, the predetermined camera maps were applied for the exposure time of the respective experiment.

3D SMLM data (STORM, PALM, DNA-PAINT) was fitted with an experimentally derived PSF model measured via z-stacks of 100 nm sized fluorescent beads as previously described^6^. For STORM data, the localizations were filtered for a lateral localization precision better than 12.7 nm, a relative log-likelihood value better than -2.9, and the first 600 frames were filtered out. For PALM data, the localizations were not further filtered. For DNA-PAINT data, the localizations were filtered for a localization precision from 0 to 12 nm and a z-coordinate of -200 nm to 100 nm. 2D DNA-PAINT data was fitted with a Gaussian PSF model and the localizations were filtered for a localization precision better than 30 nm and a PSF width of 100 to 175 nm. Diffraction-limited TIRF images were processed using the NCS software and ACsN software, respectively, as provided by the authors. As input, we use the camera maps determined via the photon-free approach described in this work, encoding the pixel-wise properties for gain, offset, and noise in case of NCS and gain and offset in case of ACsN. We parameterized the NCS MATLAB “single pixel with normalization”-algorithm by an alpha weight factor of 10, a pixel size of 0.0977 µm, an emission wavelength of 0.525 µm, a numerical aperture of 1.49, and 25 iterations. We parameterized the ACsN MATLAB app with a numerical aperture of 1.49, an emission wavelength of 525 nm and a pixel size of 108 nm, turned the video filter off and the parallel CPU option on.

### SMLM simulation

Raw 3D SMLM data were simulated in MATLAB using an experimentally derived PSF model for the microscope described above, experimentally derived camera characteristics via the photon-free approach described in this work, and photon counts parameterized by DNA-PAINT, STORM and PALM experiments described above. Camera data was simulated using a projected camera pixel width of 98 nm and the emitters were placed on the center of each camera pixel. Each emitter position was simulated for 1,000 times with the distribution of photon counts drawn from the experimentally derived distribution of the photon counts per emitter and per frame. Poisson noise was added to the photon distribution over the experimental PSF and the fluorescence signal was converted to ADU counts. The camera baseline was added, the dark current was added corresponding to the respective single frame exposure time (50 ms for PALM and STORM, 500 ms for DNA-PAINT), read noise and thermal noise was added corresponding to the respective single frame exposure time. The synthetic raw 3D SMLM data was then fitted either using a (s)CMOS-specific fitter with explicit consideration of pixel-to-pixel variations of the camera properties including dark current and thermal noise, or neglecting pixel-to-pixel variations and using the average values of the camera properties instead. The bias for each emitter position was determined as the deviation of the mean fitted coordinate from the ground truth.

For the expected root mean square error (RMSE), we followed the same approach as described above, but did not draw the photon counts from a distribution. Instead, we simulated all emitters with the same photon counts using the mean photon counts from the distribution (i.e. 3,420 photons for PALM, 9,000 photons for STORM, 35,100 photons for DNA-PAINT in case of the scientific-grade CMOS camera, and 1,900 photons for PALM, 5,000 photons for STORM, 19,500 photons for DNA-PAINT in case of the industry-grade CMOS camera). The theoretically achievable precision was calculated via the square root of the Cramér-Rao lower bound (CRLB)^20^ according to the particular PSF shape^6^.

## Data availability

Example data can be downloaded from https://rieslab.de/#accent.

## Code availability

The software for the data acquisition and analysis used in this paper is available at https://github.com/ries-lab/Accent/releases and https://github.com/jries/SMAP.

## Acknowledgements

We thank Ingmar Schoen for providing secondary antibodies for the DNA-PAINT experiments and Steve Boulant for providing the U373 cell line. We thank Alejandro Colchero for assistance in setting up the microscope, Philipp Hoess and Jervis Thevathasan for additional sample preparations, Amir Rahmani for assistance in data acquisition and Takahiro Deguchi and Sheng Liu for testing of the software. This work was supported by the European Research Council (grant no. ERC CoG-724489 to JR), the National Institutes of Health Common Fund 4D Nucleome Program (grant no. U01 EB021223 to JR), the Human Frontier Science Program (RGY0065/2017 to JR), the Engelhorn Foundation (Postdoctoral Fellowship to RD) and the European Molecular Biology Laboratory.

## Author contributions

RD conceived the project, recorded the data, analyzed the data and prepared the figures. JR supervised the project and developed the SMAP software for SMLM reconstruction and analysis. JD programmed the ACCeNT-specific plugins and scripts for Micro-Manager and Fiji. YL performed the simulations. AT prepared the live-cell and PALM samples. MK prepared the DNA-PAINT samples. UM prepared the STORM samples. RD, JR, JD and YL prepared the manuscript with input from all authors. All authors reviewed the manuscript.

## Supplementary Figures

**Supplementary Figure 1:**
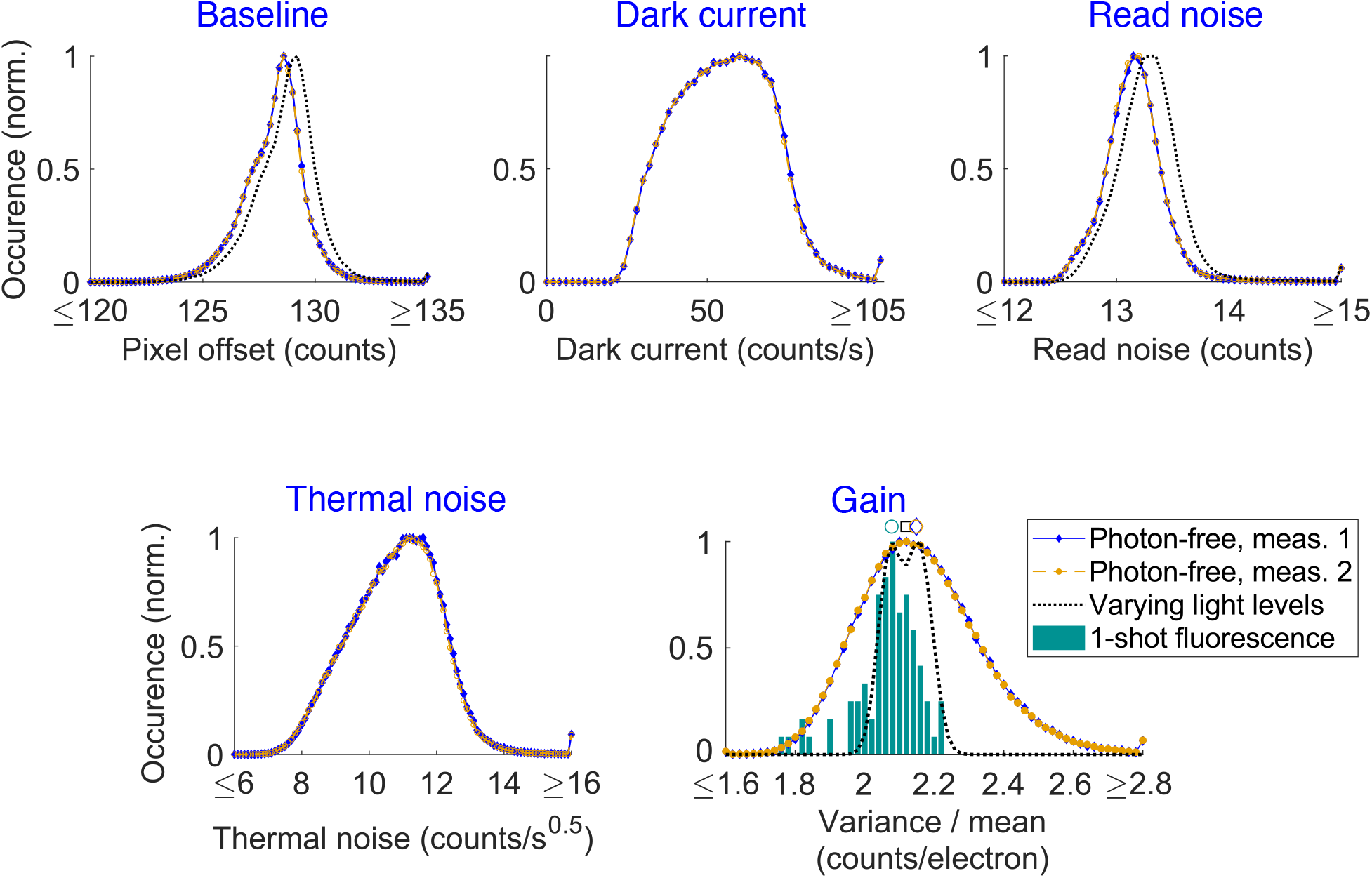
Repeated photon-free characterization of the same camera shows reproducibility of the approach. Comparison of repeated photon-free characterizations of the same camera show high similarity (blue and orange curves). For comparison: Histograms of pixel values obtained by photon-free characterization (blue curve and orange curve) and traditional characterization of using varying light levels (black dashed curve). Distribution of the gain determined via different approaches (pixelwise histogram for the photon-free and varying light levels approaches, histogram of outcomes from multiple determinations of the mean gain from the 1-shot approach). Symbols above the curves indicate the medians (blue diamond and orange circle for photon-free approach, square for vary light levels approach, circle for 1-shot fluorescence).

**Supplementary Figure 2:**
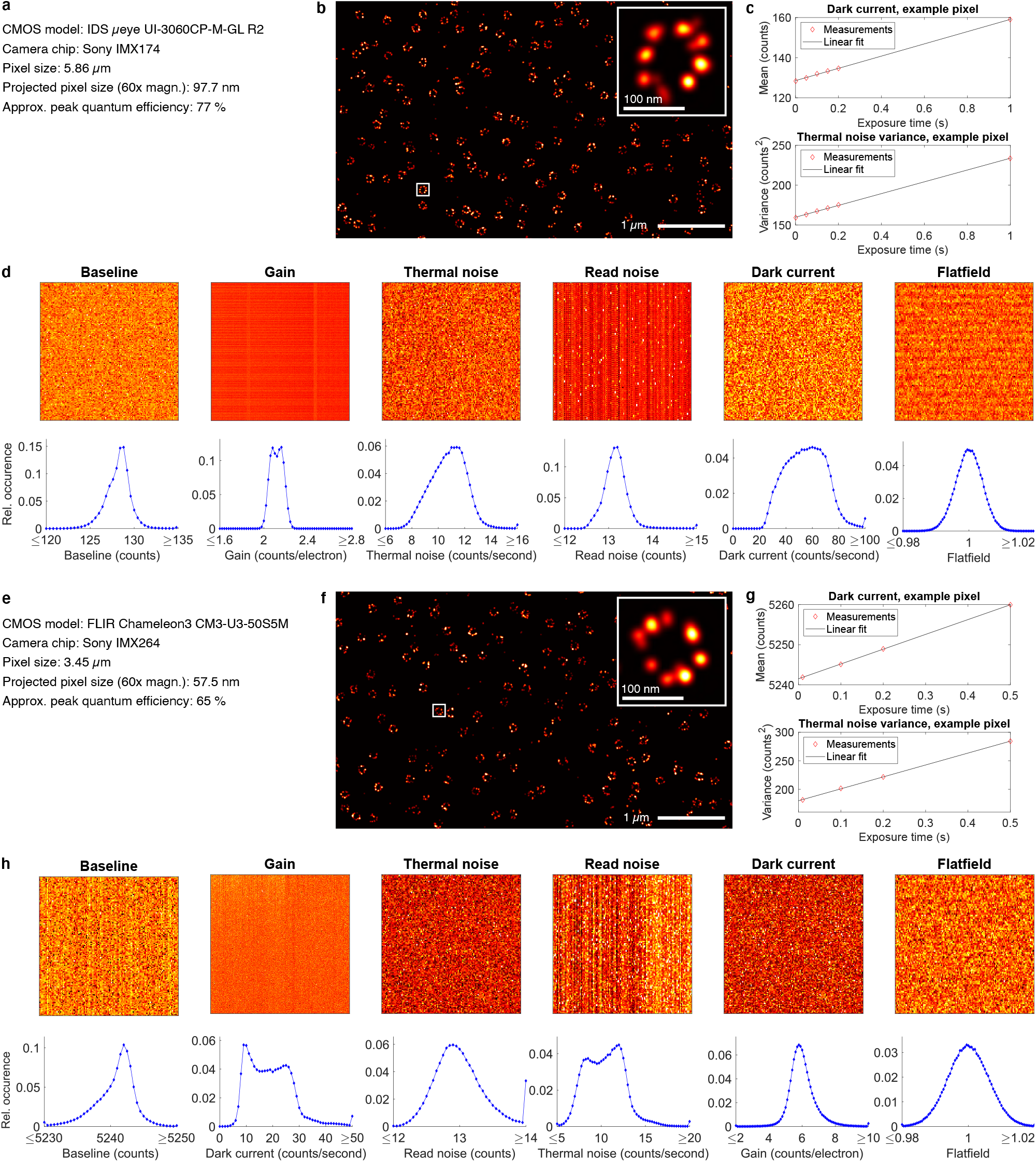
Characteristics of and STORM example images recorded by two uncooled, industry-grade CMOS cameras. **a**,**e**, Camera parameters, **b**,**f**, example STORM reconstructions, **c**,**g**, example data for exposure-time dependent signal and noise and **d**,**h**, measured camera characteristics for two different industry-grade CMOS cameras.

**Supplementary Figure 3:**
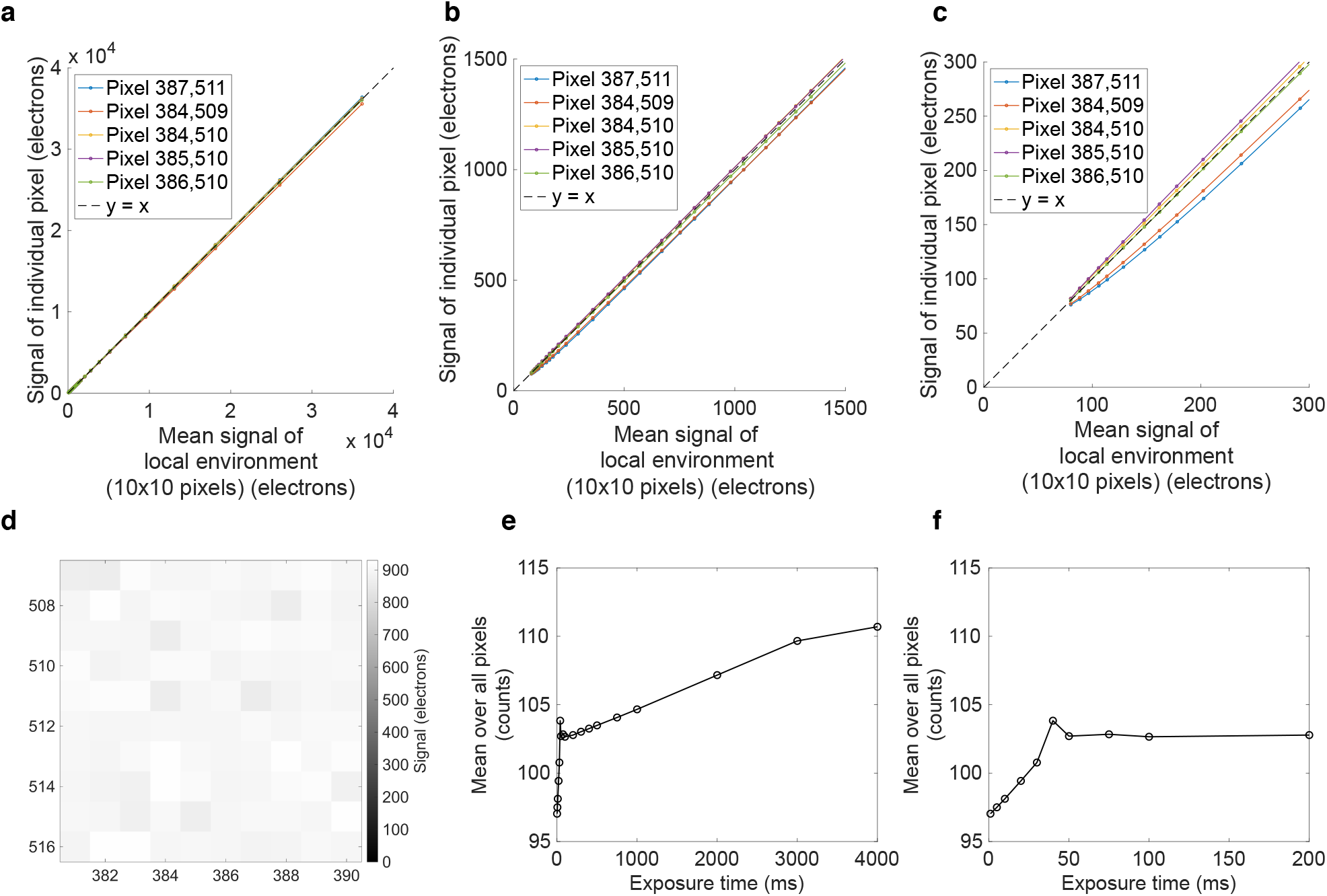
Nonlinear effects observed for a cooled, scientific-grade CMOS camera. **a-c**, We applied the traditional characterization routine of varying light levels (ensuring a spatially mostly homogeneous illumination) to our cooled, scientific-grade CMOS camera (Edge 4.2 bi, PCO) and plot the mean signal per illumination condition as a function of the mean signal of the local 10×10 pixel neighborhood for selected pixels. **a** shows 0 % to 72 % of the dynamic range of the camera, **b** shows a zoom into 0 % to 3 % of the dynamic range and c shows a zoom into 0 % to 0.06 %. While the light response of the camera is mostly linear over a large part of the dynamic range, the zoom in (c) reveals nonlinearities for some pixels particularly for about the first 200 (photo)electrons above the offset. For the DNA-PAINT experiments presented in this work, the mean background per pixel was 301 photons (and the fluorescence signal above the background was much higher), so we operate the camera in the linear regime. **d** shows the mean signal for a part of the camera. **e,f**, We recorded dark images at different exposure times and plot the mean signal of all pixels as a function of the exposure time. **e** shows the plot for 0 ms to 4,000 ms and **f** shows a zoom into 0 ms to 200 ms. The plot reveals three different regimes of dark current for the camera: The dark current appears proportional to the exposure time for 0 ms to 30 ms (corresponding to high to medium framerates often used in STORM and PALM imaging), some transitioning behavior for exposure times from 40 ms to 200 ms, and again appears proportional to the exposure time for 200 ms to 3000 ms. For the experiments presented here, we characterized the camera via the photon-free approach for exposure times from 350 ms to 3000 ms and operate the camera at 500 ms for the DNA-PAINT imaging, so the characterization as well as the experiments are performed in a linear regime.

**Supplementary Figure 4:**
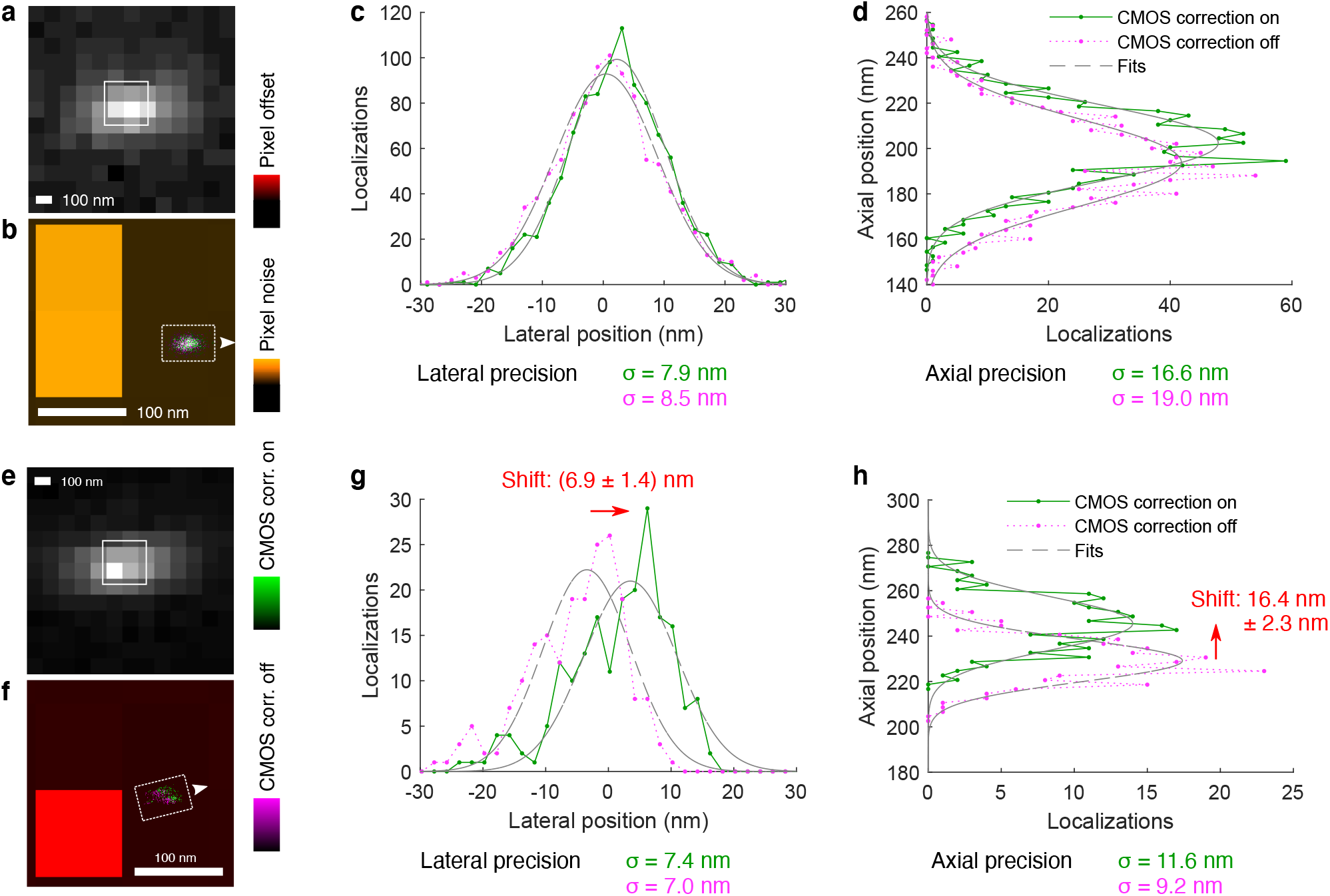
Uncorrected CMOS pixel-to-pixel variations affect localization accuracy both laterally and axially. **a**, A fluorescent bead was positioned such that its diffraction limited image was captured close to two pixels of increased noise. **b** shows a zoom into the region indicated in **a** including the pixel noise characteristics and the repeated localizations via astigmatism-based 3D PSF fitting. Magenta dots show the localizations when not using an (s)CMOS-specific fitter that takes care of individual pixel characteristics and green dots show the localizations when using an (s)CMOS-specific fitter that explicitly considers individual pixel characteristics including noise. Gaussian fits to line profiles of the localization distribution inside the boxed region indicated in **b** show that the localization precision (i.e. the standard deviation of the localizations) is increased when not using the (s)CMOS-specific fitter both laterally (**c**) and axially (**d**). **e**,**f**, Repeating the same experiment, but capturing the image of a bead in proximity to a pixel of high offset resulting from high dark current. In this case, the mean localized position is significantly shifted both laterally (**g**) and axially (**h**). The axial shift occurs because the axial position is being estimated from the shape of the PSF which becomes apparently distorted by the pixel of increased brightness when not applying a (s)CMOS-specific fitter that explicitly considers individual pixel characteristics including offset.

**Supplementary Figure 5:**
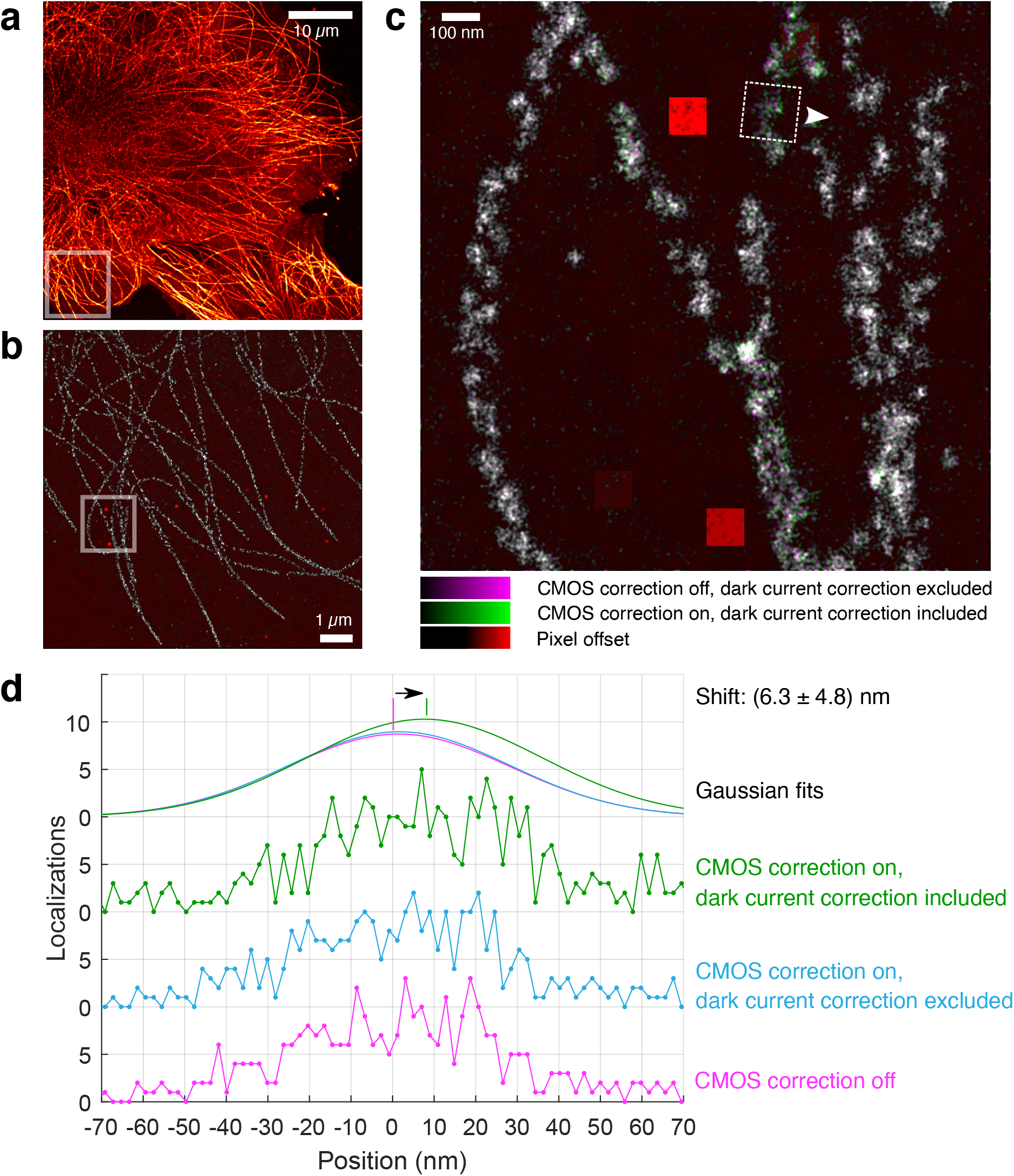
Application of CMOS specific fitting benefits from explicit correction of thermal effects. **a**, DNA-PAINT image of microtubules in a U2OS cell recorded using an uncooled, industry-grade CMOS camera. **b** shows a zoom into the region indicated by the box in **a** and **c** shows a zoom into the region indicated by the box in **b**. Localizations fitted without CMOS correction are displayed in magenta while the same localizations fitted with CMOS correction including thermal effects are displayed in green. The pixel offset, of which variations mainly stem from dark current, is displayed in red for the underlying pixel grid of the camera. **d**, line profiles according to the region indicated in **c** and Gaussian fits to determine the center position of the structure. The center becomes shifted by about 6 nm when not applying CMOS-specific fitting and, more notably, this shift persists even when the CMOS correction is applied in principle, but thermal effects are neglected.

### Supplementary Note 1: Maximum likelihood localization in the presence of sCMOS specific noise

Huang *et al*. introduced a clever way to account for pixel-specific read-noise in maximum likelihood estimation (MLE) based fitting^1^. By approximating the normal distributed readout noise (*ν*_*k*_ in the unit of electrons square, *k* denotes the pixel indices) with a Poisson distribution, one can expect the sum of the measured photoelectrons *D*_*k*_ in each pixel and the pixel-dependent readout noise *ν*_*k*_ to approximate a Poisson distribution with the mean of *u*_*k*_ + *ν*_*k*_:

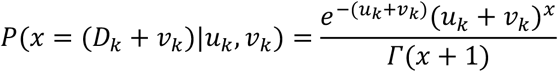

Here, *u*_*k*_ is the expected photoelectrons in pixel k calculated from the PSF model. Therefore, the conventional MLE fit for Poisson distribution can be easily applied for the pixel-dependent (s)CMOS data by adding the readout noise *ν*_*k*_ to the measured photoelectron *D*_*k*_ as the observed data and update the model function *u*_*k*_ with *u*_*k*_ + *ν*_*k*_ as the expected value. Since the exposure time-dependent thermal noise and dark current both follow the Poisson distribution and they are effectively added in the finally detected electrons, we can assume that *D*_*k*_ + *ν*_*k*_ will approximate a Poisson distribution with the mean of *u*_*k*_ + *ν*_*k*_ + *C*_*k*_ * *t* + *T*_*k*_ * *t*. Here, *C*_*k*_ denotes the dark current in the k_th_ pixel per time (electrons/second) and *T*_*k*_ denotes the variance introduced from thermal noise in the k_th_ pixel per time (electrons/second). *t* is the camera exposure time. Compared to Huang’s method considering only the effect of pixel dependent readout noise, we further take account of the effect of pixel and exposure time dependent dark current and thermal noise and incorporate them into our MLE fitting model.

### Supplementary Note 2: Software Manual Introduction

The complete ACCeNT workflow for camera characterization and SMLM fitting can be performed in two different ways:

- *EITHER* using the Micro-Manager 2 ACCeNT plugin for acquisition and simultaneous ACCeNT camera map computation (“Micro-Manager 2 calibration”) followed by SMAP SMLM fitting (“Single-molecule localization using SMAP”);
- *OR* using the Micro-Manager 1.4 ACCeNT script for acquisition followed by using the Fiji plugin for ACCeNT camera map computation (“Micro-Manager 1.4 and Fiji calibration”) followed by using SMAP for SMLM fitting (“Single-molecule localization using SMAP”).

As an alternative to the complete workflow, users can choose to perform only certain parts of the pipeline. For instance, the Micro-Manager 2 plugin allows to perform acquisition and computation independently, or the Fiji plugin can be used for computation of appropriately acquired raw camera data from other software.

To test our pipeline on provided example data, start at point 2. of the “Micro Manager 1.4 and Fiji calibration” workflow below. This includes using Fiji for ACCeNT camera calibration and using SMAP for fitting of 3D SMLM data. The example data corresponds to the 3D STORM data of nuclear pore complexes shown in Fig. 3. The example data has been recorded using an uncooled, industry-grade CMOS camera (IDS UI-3060CP-M-GL Rev.2, Sensor Sony IMX174) and can be found here: https://rieslab.de/#accent.

We successfully tested ACCeNT with the following cameras:

- UI-3060CP-M-GL Rev.2, IDS
- Edge 4.2bi, PCO
- DCC1545M, Thorlabs
- Chameleon3 CM3-U3-50S5M, FLIR
- Chameleon3 CM3-U3-31S4M-CS, FLIR
- Prime BSI, Photometrics

We did not manage to run ACCeNT with the Hamamatsu Orca Flash4.0v2 because of an apparent dark current correction implemented by the manufacturer.

#### Micro-Manager 2 calibration

1. Installation
  a. Download the latest version of Micro-Manager 2 from:
    - https://micro-manager.org/wiki/Download_Micro-Manager_Latest_Release
  b. Download the latest release of the ACCeNT Micro-Manager 2 plugin:
    - https://github.com/ries-lab/Accent/releases
  c. Place the downloaded .jar in the “mmplugins” folder of your Micro-Manager 2 installation folder.
2. Start Micro-Manager 2 using a hardware configuration that includes your camera. Refer to the Micro-Manager wiki or the Image.sc forum for any trouble regarding hardware.
  a. ACCeNT can be started from the plugins menu, under “Acquisition Tools”. The plugin consists of three steps: acquisition, processing and maps generation. All steps can be carried out together or separately.
3. Set ACCeNT parameters:
  a. Using the “…” button, select a folder in which the images will be saved.
  b. Select the name you want to give the images, the number of frames per exposure and the exposures. We advise a minimum of 15000 frames and 3 exposure times.
  c. By clicking on “Options”, you can select further acquisition options:
    - Pre-run (min): pre-acquisition run time to thermally equilibrate the camera. This is particularly necessary in the case of uncooled cameras. The total run time is an approximation as overhead exists and is camera dependent.
    - Save frames as: allows saving the frames as tiff stacks or individual images.
    - Process data: process the calibration live (in parallel) or wait for the user to start it. If the acquisition is faster than the processing, the plugin will run into a buffer overflow and crash. In such a case, run your calibration with the “separately” option.
    - Roi: sets the roi on which to perform the calibration. Roi set by Micro-Manager are ignored. If all fields are 0, then the calibration is performed on the full chip. Save the options to validate them.
4. Click on “Run” to start the acquisition. If the “in parallel” processing options was chosen, the processing also starts. The acquisition step saves the images, as well as a JSON representation of the roi (roi.roi file).
5. Once the acquisition is done, the folder and roi fields of the processing panel are updated.
  - If the processing was chosen to be in parallel to the acquisition, then it finishes soon after the last image is acquired.
  - If the processing is done separately, click on “Process”.
6. The processing step can be performed later by just loading the folder containing the images using the “…” button. The plugin automatically detects the roi.roi file and updates the fields. If no roi file is present, input the roi x, y, width and height manually.
7. The processing steps creates a calibration file (results.calb), an average and variance map for each exposure, as well as images of the various estimates present in the calibration.
8. The map generation step is automatically performed after each processing step. To run a new generation step, you can load a calibration file using the “…” button. Input the desired exposures in the exposure field and click on “Generate”. The map generation step generates the average and variance maps for the required exposures.

The calibration file (results.calb) can then be used with SMAP.

#### Micro-Manager 1.4 and Fiji calibration

In this pipeline, we make use of Micro-Manager 1.4 to generate the data. In principle the Fiji plugin can be used regardless of the way the raw data was generated. The only conditions are:

- The images are tiff (stacks or individual images)
- Each stack’s name contains “XXXms”, where XXX is the corresponding exposure time, and all stacks are in the same folder. **Alternatively**, each exposure (stack or individual images) can be in a folder with name containing “XXXms”. All exposure folders must be grouped in one folder.

1. Acquire camera frames with Micro-Manager 1.4.
  a. Download the latest version of Micro-Manager 1.4 from: https://micro-manager.org/wiki/Download_Micro-Manager_Latest_Release
  b. Start Micro-Manager 1.4 with a hardware configuration containing your camera.
  c. In “Tools”, open the “Scripting panel”.
  d. Then load “accent-acquisition_script.bsh” from the Github repository: https://github.com/ries-lab/Accent/blob/master/accent-mm1/accent-acquisition_script.bsh
  e. In the script, modify the parameters “path”, “exposures” and “numFrames”. Additionally, change the roi on line 12. The parameters of the setRoi method are in order x0, y0, width and height; with x0 and y0 defining the top-left corner of the roi. Note that certain cameras do not support this function. In such case, comment out (using “//”) line 12 and set the roi manually in Micro-Manager (refer to Micro-Manager user’s guide)
  f. Run the script to acquire the images. This can take some time, watch the console for progress and wait for the acquisition to complete.
2. **Start here to test ACCeNT on provided example data**. The example data can be downloaded at https://rieslab.de/#accent. The zip-archive for the example data contains raw data from the camera calibration, 3D stacks of beads for the point spread function (PSF) characterization and raw STORM data corresponding to Figure 3k-m.
3. Install Fiji
  a. Download Fiji: https://imagej.net/Fiji/Downloads Please note that the ACCeNT plugin uses features that have been added recently to Fiji. If you encounter errors, make sure to use the latest version of Fiji.
  b. Download the latest release of the ACCeNT Fiji plugin: https://github.com/ries-lab/Accent/releases
  c. Place the downloaded accent-fiji-1.0.jar in the “plugins” folder of your Fiji installation folder.
4. Start Fiji and the ACCeNT plugin from the plugins menu.
5. Set ACCeNT parameters:
  a. Detect the images by selecting the folder in which they are contained using the “…” button. They should appear in the table with the correct exposure time in the second column.
    - The example raw ACCeNT data is located in the folder “ACCENT_raw”
  b. Set the roi parameters x (X0), and y (Y0), e.g. used in the Micro-Manager 1.4 acquisition under point 5.
    - For the example data, keep the default values X0 = 0 and Y0 = 0.
6. Click on “Process” and wait. Processing of the example data takes about 20 minutes (Windows 64-bit operating system, i5-4690K CPU @ 4×3.5GHz, 32 GB memory).
7. Output files:
  a. The processing steps creates a calibration file (results.calb) to be used with SMAP, as well as different camera maps (in units of ADU counts unless stated otherwise):
  b. The mean pixel values for each exposure time are saved as Avg_XXXms.tiff, where XXX denotes the exposure time in ms.
  c. The variance pixel values for each exposure time are saved as Var_XXXms.tif, where XXX denotes the exposure time in ms.
  d. The computed baseline is saved as Baseline.tiff.
  e. The computed dark current per 1 second is saved as DC-per_sec.tiff.
  f. The square of the computed read noise (in units of ADU counts squared) is saved as RN_sq.tiff.
  g. The square of the thermal noise per 1 second (in units of ADU counts squared) is saved as TN_sq_per_sec.tiff.
  h. The computed gain (in units of ADU counts per electron) is saved as Gain.tiff.
  i. The computed offset maps for arbitrary exposure times (default values are 15 ms, 20 ms, 30 ms, 50 ms and 100 ms) are saved as generated_Avg_XXXms.tiff, where XXX denotes the exposure time in ms.
  j. The computed variance maps for arbitrary exposure times (default values are 15 ms, 20 ms, 30 ms, 50 ms and 100 ms) are saved as generated_Var_XXXms.tiff, where XXX denotes the exposure time in ms.
  k. The R_sq_YYY.tiff files show the R^2^ values for inspection of the quality of the fits used to compute baseline (R_sq_avg.tiff), read noise squared (R_sq_var.tiff) and gain (R_sq_gain.tiff).
    - You can inspect the generated camera maps of the example data using Fiji. For instance, open the “DC_per_sec.tiff” in Fiji. While the mean dark current is 54 counts/s (using Analyze -> Measure; set mean via Results -> Set Measurement), you will find that pixel (256,289) features significantly higher dark current of 1429 counts/s (placing the cursor over the pixel reading the value from the main Fiji window). Visual inspection of the image shows other pixels of pronounced dark current, e.g. (220,315), (289,284) and (266,247). To find the dark current in units of electrons/s, open the “Gain.tiff” file and measure the median gain value (using Analyze -> Measure; set median via Results -> Set Measurement). For the example data, the gain is 2.16 counts/electron. Hence, the mean dark current is (54 counts/s) / (2.16 counts/electron) = 25 electrons/s.
    - To inspect the read noise, open the “RN_sq.tiff” and calculate its square root (using Process -> Math -> Square Root). The mean read noise is 13.2 counts, corresponding to (13.2 counts) / (2.16 counts/electron) = 6.1 electrons.
8. Though not needed for fitting of single molecule localization data in SMAP, ACCeNT can create exposure time specific camera maps. These can for instance be used to explore camera characteristics, find particularly bad pixels, or as input for software other than SMAP. The map generation step is automatically performed after each processing step. To run a new generation step, you can load a calibration file using the “…” button. Input the desired exposures in the exposure field and click on “Generate”. The map generation step generates the average and variance maps for the required exposures.
  - For the example data, type “999” into the field “Exposures (ms):” and click “Generate”. The generated camera maps for offset and variance corresponding to 999 ms exposure time will be saved to the folder “ACCENT_raw”. To check the consistency with the measurement, open both the “Avg_999ms.tiff” (the mean pixel values directly from the measured data) and the “generated_Avg_999ms.tiff” (the computed offset values for this exposure time) files in Fiji. Compute the difference (using Process -> Image calculator…; Image 1: Avg_999ms.tiff, Operation: Subtract, Image 2: generated_Avg_999ms.tiff, check Create new window, check 32-bit (float) result). The mean difference between the measured and computed maps is –0.042 counts corresponding to (−0.042 counts) / (2.16 counts/electron) = -0.019 electrons.

#### Single-molecule localization using SMAP

1. Install SMAP from: https://github.com/jries/SMAP. The version from time of publication is available at: https://github.com/jries/SMAP/releases/tag/v210404. following the instructions. Alternatively, a stand-alone version for PC and Mac can be downloaded at: https://rieslab.de/#software. Please follow the installation instructions provided with the executables. Familiarize yourself with SMAP by consulting the documentation, using example data downloaded at https://rieslab.de/#accent. Make sure to install Micro-Manager and select its path in the Preferences menu.
2. Copy the ‘results.calb’ file in the SMAP/settings/cameras directory.
3. As described in the user guide (<u>SMAP_UserGuide.pdf(embl.de)</u>, page 6), add your camera to the camera manager. In the ‘correctionfile’ field, select the results.calb file. For the example data, the camera is already registered in the camera manager.
4. For fitting of 3D SMLM data, perform the 3D calibration as described in the SMAP user guide (https://www.embl.de/download/ries/Documentation/SMAP_UserGuide.pdf, page 9 “experimental PSF model”). For the provided example data, you can EITHER skip this step and use the file: “60xOil_sampleHolderInv CC0.140_1_MMStack.ome_3dcal.mat” from the “PSF_raw” folder in the second next step, OR perform the 3D PSF calibration yourself, based on 3D stacks of 100 nm Tetraspeck beads:
  - Open “Plugins -> Analyze -> sr3D -> calibrate3DsplinePSF”
  - On the new window, click “Run”
  - On the new window, click “Select camera files”
  - On the new window, click “add dir”
  - Navigate to the “PSF_raw” folder and mark all folders
  - Click “Open”
  - In the previous window, click “Done”
  - In the previous window, click “Calculate bead calibration” and wait. Processing of the example data takes about 2 minutes (Windows 64-bit operating system, i5-4690K CPU @ 4×3.5GHz, 32 GB memory, GeForce GTX 970 GPU).
5. In the SMAP main window select the ‘Localize’ tab and the ‘Input Image’ subtab to load the raw SMLM data via “load images”.
6. For the example data, open the first Tiff-file in the “SMLM_raw folder”. Click “set Cam Parameters” to validate that the camera has been recognized. Do not change any values and click “OK”. The example data corresponds to the 3D STORM data of nuclear pore complexes shown in Fig. 3k-m. The example data has been recorded using an uncooled, industry-grade CMOS camera (IDS UI-3060CP-M-GL Rev.2, Sensor Sony IMX174). It contains 19000 frames. The recording was manually stopped after bleaching of most fluorophores. Check ‘correct offset’.
7. In the SMAP main window, change to the ‘Fitter’ subtab, select ‘Spline’ as the fitting model via the drop-down menu, click “Load 3D cal” and load the 3D PSF calibration file. For the example data, load the file: “60xOil_sampleHolderInv CC0.140_1_MMStack.ome_3dcal.mat”.
8. Check “sCMOS”.
9. Check “RI mismatch”
10. Click “Preview” to test fitting.
11. If successful (i.e. you get the message “preview done”), press “Localize” to start the localization process. For the provided example data, this takes about 6 minutes (Windows 64-bit operating system, i5-4690K CPU @ 4×3.5GHz, 32 GB memory, GeForce GTX 970 GPU). The file containing the localizations is saved as “SMLM_raw_sml.mat” in the main folder.
12. You can repeat the fitting process by unchecking the “sCMOS” check box in the “Fitter” tab, unchecking the “Correct offset” checkbox in the “Input Image” tab and compare the results.

For the provided example data, go to the “Input Image” tab and unclick “Correct offset”, go to the “Fitter” tab and unclick “sCMOS”, click “Localize” and wait. This will take another about 6 minutes.

- In the main window, click “Load” and navigate to the main folder of the example data. Mark both “SMLM_raw_sml.mat” (the fitting results using the CMOS fitter with the ACCeNT calibration) and “SMLM_raw_2_sml.mat” (the fitting result not using the CMOS fitter). Click “Open”. If you have not performed the fitting step yourself, you can alternatively load the provided localization files “provided_SMLM_CMOS_fitter_sml.mat” and “provided_SMLM_no_correction_sml.mat”.
- Change to the “Render” tab.
- In the drop-down menu, select “ SMLM_raw_sml”
- Change “LUT:” to “green” via the drop-down menu.
- Next to “quantile”, change the value to “–3.0”
- Set the localization filters on the right to the following intervals (lower bound is left value, upper bound is right value):
  - locprec: [0, 30]
  - z: [-200, 300]
  - frame: [600, Inf]
- Next to “Layer1”, click “+”.
- In the drop-down menu for the new “Layer2”, select “ SMLM_raw_2_sml”
- Click the button “frame” to re-activate the frame filter.
- Click “Inv” next to “LUT:”.
- Click “Render”.
- In the “format” region of the main window, click “Reset”.
- To perform drift correction, change to the “Process” tab, check “Correct z-drift” and click “Run.” Drift correction will take about 2 minutes.
- Go back to the “Render” tab and click “Render”.
- In the “format” region of the main window, change the value of Pixrec (nm) to “3” and press enter.

You can now inspect the reconstructed STORM image. Move around by right-clicking the region you want to bring to the center of the ROI. Data that has been fitted with the CMOS fitter is shown in green, data that has been fitted without the CMOS fitter is shown in magenta. Accordingly, regions where localizations from both fitting processes coincide well are displayed greyish.

To visualize the reconstructed STORM image in 3D, in the “layers” part of the main window, uncheck the box of “2”. Go to “Layer1”, change “Colormode:” to “z”, change “LUT:” to “jet”, change the values next to “c range” to –100. In the “format” region of the main window, change the value of “Pixrec (nm)” to 10, press enter and click “Render”. The z-coordinate of the dataset that has been fitted using the CMOS-specific fitter is now color-coded. Move around by right-clicking the region you want to bring to the center of the ROI.

## Supplementary Videos

**Supplementary Video 1**: 3D reconstruction of the four nuclear pore complexes shown in Figure 2m.

**Supplementary Video 2**: Time-lapse live-cell TIRF data of GFP-tagged AP2 in U373 cells recorded using an uncooled, industry-grade CMOS camera. Left: Unprocessed data directly from the camera, Center: NCS-processed data after ACCeNT-calibration of the camera, Right: ACsN-processed data after ACCeNT-calibration of the camera. Note that for the ACsN algorithm, the parameter for the projected pixel width was chosen as 108 nm (instead of 98 nm which corresponds to the physically correct value) since the correct value led to a considerable loss in resolution.

